# The migration of an expanding sea turtle population alters the structure of marine megafauna communities

**DOI:** 10.1101/2025.09.17.676690

**Authors:** Stuart Robert Brian Negus, Elton Neves, Aírton Jesus Lima, Carolina Oujo, María E. Medina Suárez, Sandra M. Correia, Anaise Andrade, Ivanildo Freitas, Thomas Reischig, Kenydjeer Lima Rodrigues, Christophe Eizaguirre, Gail Schofield

## Abstract

Interactions shape community structures, yet the impact of large migratory influxes on communities, particularly marine ones, remains elusive. Using drone surveys, we investigated changes of marine megafauna communities (e.g., sharks, rays), during the mass migration of loggerhead sea turtles (*Caretta caretta*) in Cabo Verde, one of the largest, and growing, rookeries globally and a biodiversity hotspot. In high nesting density areas, with over 6,000 turtles detected in-water at the season’s peak, the community structured with sharks distributing between the turtles and the coastline. In low-density areas (∼500 turtles), no structuring was detected. After the season’s peak, increased shark abundance and shoreward movement suggest they anticipate a nutrient pulse from emerging hatchlings. Our results highlight the role of sea turtle abundance in structuring marine communities, both as a high-quality food resource (hatchlings) and a likely non-consumable species (adults). Overall, we show the significant impact of migratory species on marine megafauna communities.

## Introduction

Mass animal migration involving thousands of individuals moving between habitats and communities is a common life history strategy across taxa to reach foraging, breeding, or nesting areas (Shaw 2016). Individuals may migrate as part of their ontogeny or seasonally due to resource fluctuations or environmental change (Dingle 2014). By occupying favourable habitats at certain times and life stages, migratory individuals increase survival, reproduction and ultimately their Darwinian fitness (Dingle 2014).

In terrestrial and freshwater ecosystems, migratory species can alter the trophic structure of the communities they enter (e.g., mammals (Anderwald *et al.* 2015), insects (Chapman *et al.* 2015), birds and fish (Brönmark *et al.* 2014)), by displacing resident species (Bauer & Hoye 2014). This results in localised competition that increases interspecific interactions and selection pressures (Bauer & Hoye 2014; Bellard *et al.* 2012; Dingle 2014; Schlägel *et al.* 2020). For example, red deer migrations (*Cervus elaphus*) negatively impact resident chamois (*Rupicapra rupicapra*) and ibex (*Capra ibex*) populations as they occupy similar dietary niches. As a result, the large migratory deer populations push chamois populations outside of their preferred areas (Anderwald *et al.* 2015).

Abundant migrants also change trophic dynamics by providing resource pulses, attracting predators through migratory coupling or increasing ecosystem productivity (Bauer & Hoye 2014; Giroux *et al.* 2012; Hulbert *et al.* 2005; Ostfeld & Keesing 2000). Individuals may migrate to resource pulse areas to either feed on the migratory species (e.g., migratory coupling) or benefit indirectly; e.g., in marine ecosystems the transport of nutrients between communities can lead to plankton blooms and subsequent foraging (Furey *et al.* 2018; Hulbert *et al.* 2005; Levi *et al.* 2015; Shaw 2016). Migration hence can supplement local food webs, increasing consumer growth and reproduction (Giroux *et al.* 2012).

For coastal and marine megafauna, data on community-scale dynamics are scarce (Grémillet *et al.* 2022). While tracking individuals through telemetry has helped to understand how species interact, this is often limited to a few individuals and may not represent the whole community (Bro-Jørgensen *et al.* 2019; Costa-Pereira *et al.* 2022). To overcome those limitations, a complementary alternative exists in the form of unoccupied aircraft systems (hereafter, aerial drones). Aerial drones overcome the limitations of tracking, as they can be used to assess marine assemblages (Kelaher *et al.* 2020), resource competition (Schofield *et al.* 2022), and spatial community structure (Dickson *et al.* 2022) by capturing regional, fine-scale data on hundreds of individuals simultaneously (Dickson *et al.* 2022). Aerial drones hence have the potential to assess whether and how the migration (arrival and departure) of marine species in large amounts affect the structure of a resident community, composed, among others, of predators of the migratory species.

Sea turtles migrate seasonally from foraging to nesting grounds (Godley *et al.*, 2010; Hays & Hawkes, 2018), sometimes at large scales, impacting the dynamics of the local species at their coastal nesting grounds. Turtles transport nutrients between habitats and provide prey either as eggs/hatchlings or even as adults (Bjorndal & Jackson 2003; Heithaus *et al.* 2005, 2008). Hatchlings are eaten by sharks (Fitzpatrick *et al.* 2012), rays and large fish (Louro *et al.* 2022) whereas larger sharks (e.g. tiger shark, *Galeocerdo cuvier*) can consume adults (Heithaus *et al.* 2005, 2008). The mass arrival of adult turtles and hatchlings could result in a resource pulses for predators, potentially changing how predators distribute themselves near nesting sites and even potential spatially restructuring entire communities (Furey *et al.* 2018). It could also result in competition for space in the coastal areas with non-predator species. However, whether the mass arrival of sea turtles does change marine community structure remains unknown, as do the possible thresholds that elicit those changes if they exist.

Here, we determined the spatial community structure of marine megafauna detected by aerial drones on Boa Vista island, Cabo Verde. This rookery is likely the world’s largest for nesting loggerhead turtles (*Caretta caretta*), where over thirty thousand turtles nest yearly (Taxonera *et al.*, 2022). Boa Vista island hosts about 60% of the total nesting population and is a biodiversity hotspot, with communities that contain elasmobranchs (Rosa *et al.* 2023), cetaceans (Wenzel *et al.* 2020) and large fish species (Duarte & Romeiras 2009). Using this biodiversity hotspot and the natural experimental setup emerging from the geographic differences in nesting distributions, we evaluated whether the community dynamics change throughout the nesting season when the mass arrival of turtles into high- and low-nesting density areas occurs. We evaluated the in-water species abundance, and distributions, as well as the distance between turtles and sharks, and the distance between the different species to the shoreline. We hypothesised that: i) communities will change as the nesting season progresses, with the most significant changes being detected at the peak of the nesting season where most migratory turtles have arrived near nesting sites, ii) predatory sharks will distribute close to the shoreline to increase predation on hatchlings late in the nesting season at the start of the hatchling season, and iii) spatial community structure in the low nesting density area may remain relatively unchanged, as the lower number of turtles found there may not be sufficient to alter community dynamics and interactions.

## Materials & Methods

### Aerial Drone Surveys, Data Collection and Processing

This study was conducted on Boa Vista island, Cabo Verde (N: 16.095011, E: -22.8070835) in the North East Atlantic during the loggerhead turtle nesting season (June – October) and during hatchling emergence (August – end October for this study, Figure 1a).

**Figure 1.**
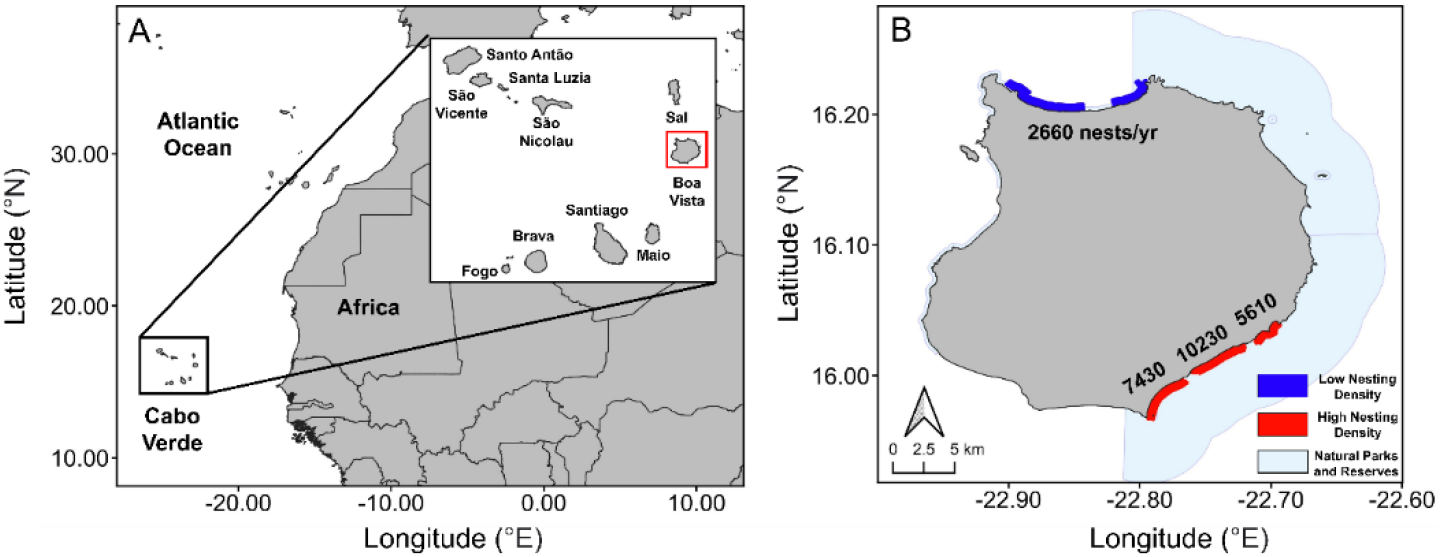
**(A)** Boa Vista in Cabo Verde, in the East Atlantic. (**B**) Locations of aerial drone surveys and the average loggerhead turtle nest numbers between 2018 and 2021 (blue for the low-density area, red for the high-density area). Nature reserves and parks that extend out into the ocean are shown as light blue polygons.

Pilot aerial drone surveys were conducted along 46 km of the coastline of Boa Vista Island before focusing on key regions (Figure 1b) on a biweekly basis. We used a DJI Phantom 3 Professional^TM^ and a DJI Mavic 2 Zoom^TM^ drone (Shenzhen, China; https://www.dji.com; accessed 14 October 2022) flown at 60 m and 70 m altitude respectively, for a 100 m field of view. This altitude enabled exploring large-scale distributions, while maintaining taxonomic identification abilities (Dickson et al., 2022a, 2022b). Surveys covered 1 to 4.5 km of offshore transects separated by 100 m up to 600 m offshore (Figure 1b). We manually reviewed footage, recording detected marine megafauna species’ locations (Dujon *et al.* 2021). The identification of species was determined using body features (e.g., carapace length/width ratio) and swimming patterns (e.g., propelled, flapping; Table 1; (Hart *et al.* 2014; Stokes *et al.* 2023). We identified 70% of sea turtles, 41% of sharks, and 2% of rays to a species level (Table 1). As not all individuals could be identified at the species level, they were then grouped into upper taxonomic groups (i.e., sea turtles, sharks, fish, and rays). All statistical analyses, unless specified, were performed in R version 4.2.2 (R Core Team 2022).

### Population trends in space and time

To evaluate the impact of the influx of loggerhead turtles on communities in this near-shore area, we counted the total number of individuals for each marine megafauna group at two-week intervals throughout the nesting season. In total, surveys were conducted at 13 individual sites within five regions across Boa Vista island: north (n = 3 sites), east (n = 4), southeast (n= 3), south (n = 1) and west (n = 2, Figure 1b; Figure S1). The southeast hosts the highest nest densities (Marco et al., 2011; Louro et al., 2022), so we analysed it separately. We counted individuals per taxon for each region at two-week intervals (Total transects = 482) and standardized this by the number of surveys flown within that interval.

Based on the population dynamics detected for sea turtles, we defined the nesting season as the succession of four periods: the first period represents the start of the nesting season when turtle counts are low (1^st^ June – 14^th^ July) and was used as the baseline to test against the changes over time. Then, we defined the second period as the increase period, when turtle abundance rises until a peak is reached (15^th^ July – 14^th^ August). The third period is defined as the decrease period when turtle abundance begins to decline from the peak (15^th^ August – 30^th^ September). The end of the season was the final period which is defined as when turtle numbers remain consistently low after the decline (1^st^ October – 31^st^ October). We hypothesized the community composition and spatial structure changes are most likely at the peak nesting season and that once nesting turtle abundance has decreased and remains low, the community composition and spatial structure will cycle back to the early conditions.

### Community trends between regions and during the nesting season

To determine the changes in the marine community composition of Boa Vista, abundance data of each taxonomic group was log+1 transformed. A Bray-Curtis dissimilarity matrix from the R package *vegan* was calculated on the log+1 transformed abundances (Oksanen *et al.* 2022). An ADONIS (Analysis of variance using Distance Matrices) was then used to test how marine community compositions changed in relation to the four described periods and nesting density area (low nesting density in the north vs high nesting density in the southeast, with the sites within each geographic area as replicates) as an interaction. Further pairwise ADONIS tests were performed, using Bonferroni correction, to determine where the differences were found across the four periods and nesting densities. A SIMPER (Similarity Percentage) analysis was then performed to determine what taxonomic group explained best the dissimilarities among spatial and temporal communities. Both ADONIS and SIMPER analyses were reperformed on the abundance data of marine megafauna, but with sea turtles removed, to causally link sea turtle abundance to changes in community structure.

### Spatial structure and distributions

We evaluated space used by turtles for each region using the 95% kernel density estimates (KDEs) for home ranges with the *adehabitatHR* package (Calenge 2006). To this end, we used the GPS data of all turtles detected to calculate turtle space use for each region. We then calculated the overlap between sharks, fish, and rays (hereafter, referred to as the resident community) with sea turtle space use, using binomial generalized linear models (GLMs) with the period of the season and the region as fixed single and interaction factors respectively. We defined the spatial community structure as the proportion of overlap between the resident community and turtle space use, with 1 showing the community is not structured (i.e., no separation of space use by taxa), and 0 for structured communities (i.e., similar taxa using specific areas).

### Distance from the shoreline and between marine megafauna groups

To examine if a change in the satial community structures occurred due to predator-prey or turtle density-dependent dynamics, we determined nearest-neighbour distances between and within marine megafauna groups. Nearest-neighbour distances were calculated within the average area surveyed by the drones (3934.5 m^2^) by survey day. Using linear-mixed effects models (LMEMs), we calculated the relationships between average nearest-neighbour distances and nesting density area, the periods of the nesting season and their interactions. Because we surveyed multiple sites within high- and low-density areas, those sites were included as random factors. We removed any taxa that were not represented at every stage of the nesting season or not found at both the high- and low-density areas (e.g., oceanic and coastal rays were not detected in transects at the start of the nesting season).

To quantify if species distributions correlated with the densities of turtles, we calculated the distance of each individual animal to coastline in metres using QGIS version 3.28.3. We used an LMEM to test whether the average distance varies among taxonomic groups, nesting density, and the periods of the nesting season as well as their interactions. Sites were included as random factors.

## Results

A total of 482 aerial drone transects were flown from 9^th^ June – 19^th^ October 2021. We detected 14,773 individual animals, of which 95% were sea turtles (n = 14,139). The other taxonomic groups that were recorded were sharks (n = 311), individual large fish (n = 144), oceanic rays (n = 63), fish shoals (n = 59) and coastal rays (n = 57).

### Population and community trends across regions and the nesting season

Sea turtles were first detected at the nesting grounds between June 1^st^ and July 14^th^ (southeast mean number of turtles per survey: 4 turtles per survey, north: 2 turtles per survey). The major increase occurred between July 15^th^ and August 14^th^ (southeast = 226 turtles per survey, north = 15 turtles per survey; Figure 2, Figure S2). A gradual decline was detected on August 15^th^ and September 30^th^ (southeast = 183 turtles per survey, north = 10 turtles per survey). Ultimately, the end of the nesting season began from October 1^st^, when almost no turtles were detected (southeast = 8 turtles per survey, north = 2 turtles per survey; Figure 2). As per the definition of the nesting season periods, the community composition changed between those four stages (ADONIS: F_*_= 9.66, *p* < 0.001; Figure 2).

**Figure 2.**
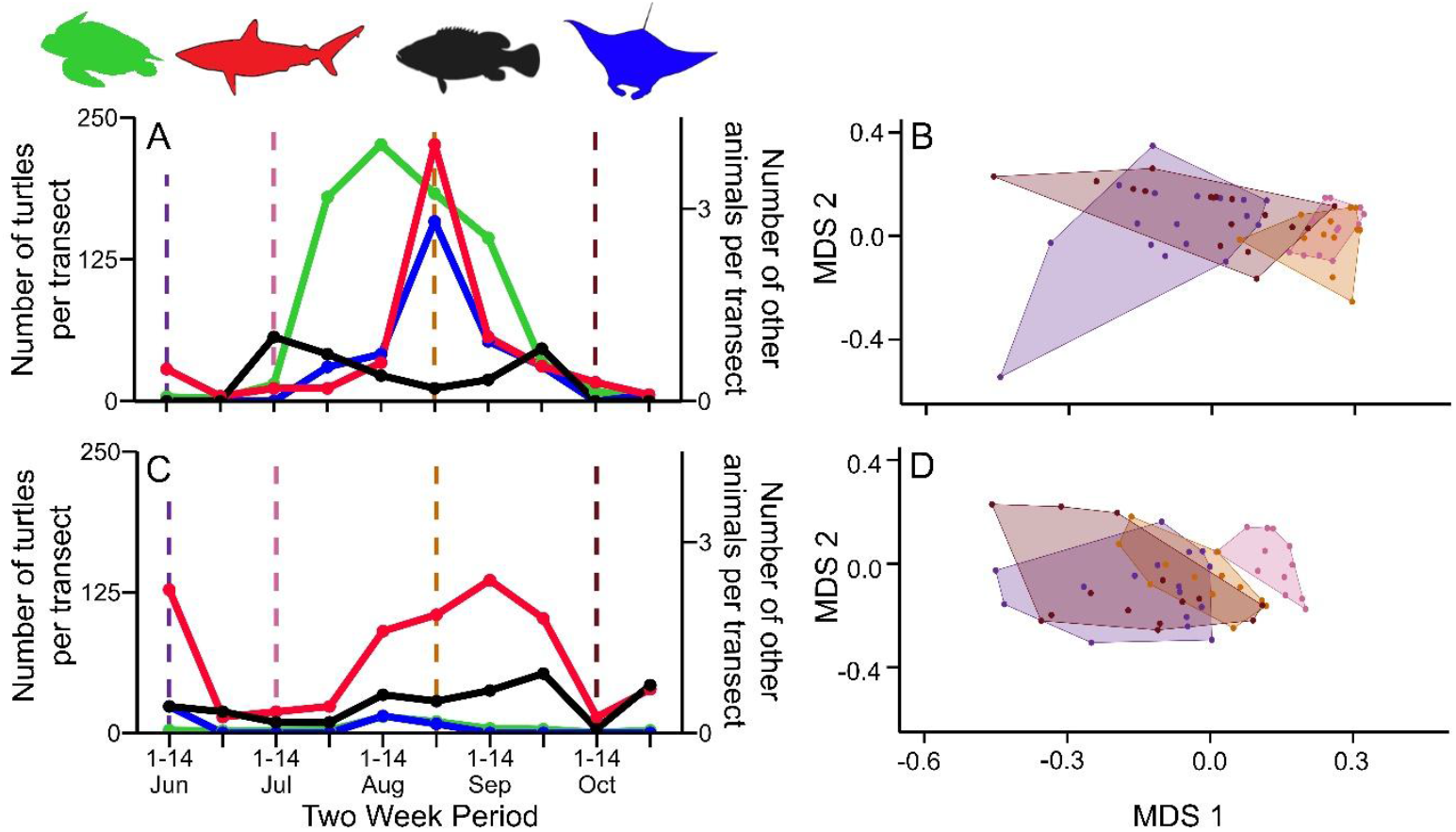
Mean number of turtles per transect (left axis) and other marine megafauna species (right axis) and non-dimensional scaling plots of marine communities in high-density (southeast: **A-B**) and low-density (north: **C-D**) nesting areas. Abundances in **A** and **C** are given as sea turtles (green), sharks (red), oceanic and coastal rays (blue) and individual fish and fish shoals (black). Dashed lines in **A** and **C** and polygons in **B** and **D** are the periods of the nesting season, given as the start of the nesting season (purple), increase of sea turtle numbers (pink), decrease in sea turtle numbers (orange) and the end of the season (brown). For **B & D**, refer to Table S2-4 for SIMPER analysis results.

Specifically, in the high nesting density area, shark and ray numbers were generally low but at their highest between August 15th and August 31st (4 sharks; 3 rays; Figure 2a). Marine megafauna communities in the high-density area were significantly different between the start of the season and the increase period (ADONIS: *p* = 0.028). No differences were detected between the increase and decrease periods (ADONIS: *p* = 1, Figure 2b), suggesting a mirrored dynamic between the arrival and departure of turtles. The communities in the decrease period were significantly different to those at the end of the season (ADONIS: *p* = 0.028), but the end of the season communities did not differ from those at the start of the season (ADONIS: *p* = 1, Figure 2b). This suggests that communities return to a composition similar to that when turtle abundance is lowest, displaying a cyclic dynamics.

In the low nesting density area, ray and shark abundances were comparable to the abundance of those groups in the high-density area (Figure 2c). Comparing community composition across the periods of the nesting seasons can show whether changes still occur despite fewer turtles. Marine megafauna communities in the low-density area were different between the start of the season to those at the increase period (ADONIS: *p* = 0.028; Figure 2d). The community composition at the increase period was also different to that at the decrease period (ADONIS: *p* = 0.028; Figure 2d). However, the communities detected during the decrease period were not significantly different to those either at the start (ADONIS: *p* = 1) or the end of the season (ADONIS: *p* = 0.504; Figure 2d). This result suggests that, like the high-density area, a cyclic dynamic in species composition occurs in the low nesting density area, but the cycle returns to the baseline sooner in the season.

When repeating tests without sea turtles, which form the core of the community during the peak nesting season, we also detected a change in the community composition throughout the four periods of the nesting season, and at different nesting density areas (ADONIS: F_*_= 4.204, *p* < 0.001). However, pairwise tests revealed that no differences were found at high nesting density areas throughout the nesting season (ADONIS: start of season-increase: *p* = 0.224, increase-decrease: *p* = 1, decrease-end: *p* = 1) or at the low nesting density areas (ADONIS: start of season-increase: *p* = 1, increase-decrease: *p* = 1, decrease-end: *p* = 1). This result shows that subtle seasonal changes in community composition occur throughout the nesting season, but these changes are amplified by sea turtles.

### Spatial community structure across the nesting season

The overlap between the resident community and the turtle space use changed during the nesting season and differed by density areas (GLM: *χ*^2^ = 28.93, *p* < 0.001; Figure 3, Figure S4.3-S4). In the high nesting density area, the overlap between the resident community and turtle space use did not change throughout the season (mean proportion of overlap ± standard error, the start of the season: 0.78 ± 0.10 community n = 18, increase: 0.84 ± 0.05 n = 49, decrease: 0.60 ± 0.09 n = 129, end of season: 0.75 ± 0.07, all *p* > 0.05; Figure 3a, c, Figure S3-S4). In the low nesting density area, however, at the end of the season (0.63 ± 0.06, n = 70) the resident community overlapped less with turtle space use compared to the start of the season (0.98 ± 0.02, n = 49, *p* = 0.03), the increase (0.96 ± 0.03, n = 48, *p* = 0.01) and the decrease periods (0.98 ± 0.02, n = 58, *p* = 0.02, Figure 3b, d, Figure S3-S4). These results suggest that the local communities in the high nesting density area share space with turtles and that a cryptic spatial structure may exist.

**Figure 3.**
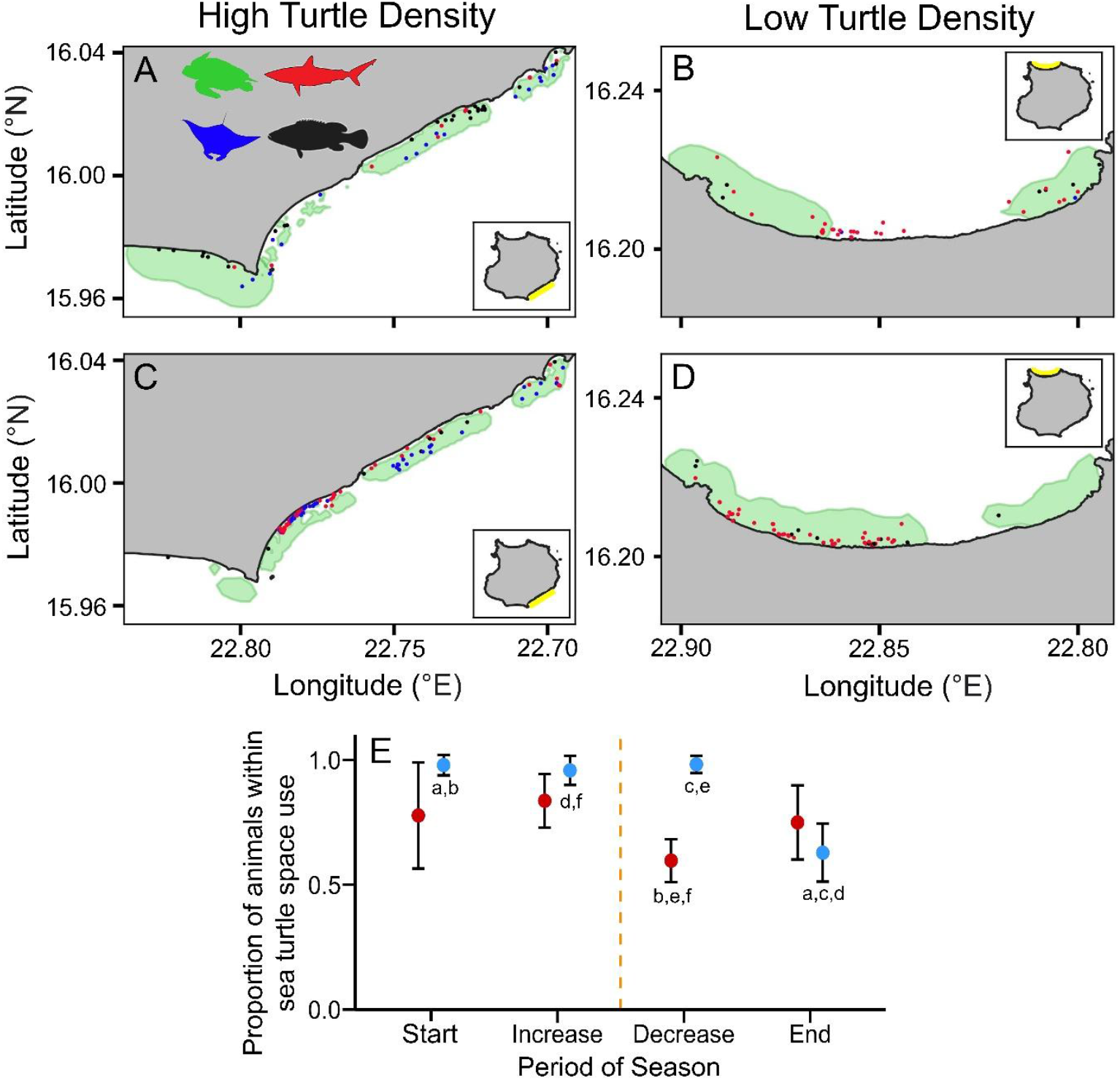
Turtle space use (95% Kernel density, green polygons) and overlapping resident community during the (**A-B**) increase to peak turtle numbers and (**C-D**) decrease from the peak turtle numbers in the high-density (left-side maps, southeast Boa Vista) and low-density (right-side maps, north Boa Vista) areas. Marine megafauna groups included sharks (red points), rays (blue), and fish (black). Matching letters highlight significant differences. (**E**) The average proportion of marine megafauna groups that were within sea turtle space use in high-density (red) and low-density (blue) areas throughout the nesting season (mean ± standard error).

### Cryptic community structure

To determine whether there is a cryptic spatial community structure in the high nesting density area, we calculated the distance between sharks and turtles. In general, as expected, turtles were closer to sharks in the high-density area than in the low-density area (Figure 4a, b) likely because of limited shallow areas. Turtles and sharks were detected closer to each other as the season progressed (log mean ± standard error, start of season: 4.51 ± 0.36 n = 9, increase:3.03 ± 0.19 n = 12, decrease: 3.33 ± 0.11 n = 74) and the distance between them increased again as the nesting season progress towards its end (4.66 ± 0.79 n = 16, all *p* = 0.05, Figure 4a). This suggests that both groups occupy similar areas that become highly populated when turtle abundance is high. In the low-density area, sharks and turtles were at their closest in the increase period (4.12 ± 0.21, *p* < 0.001; Figure 4b), as expected with some overlapping but in less populated areas.

**Figure 4.**
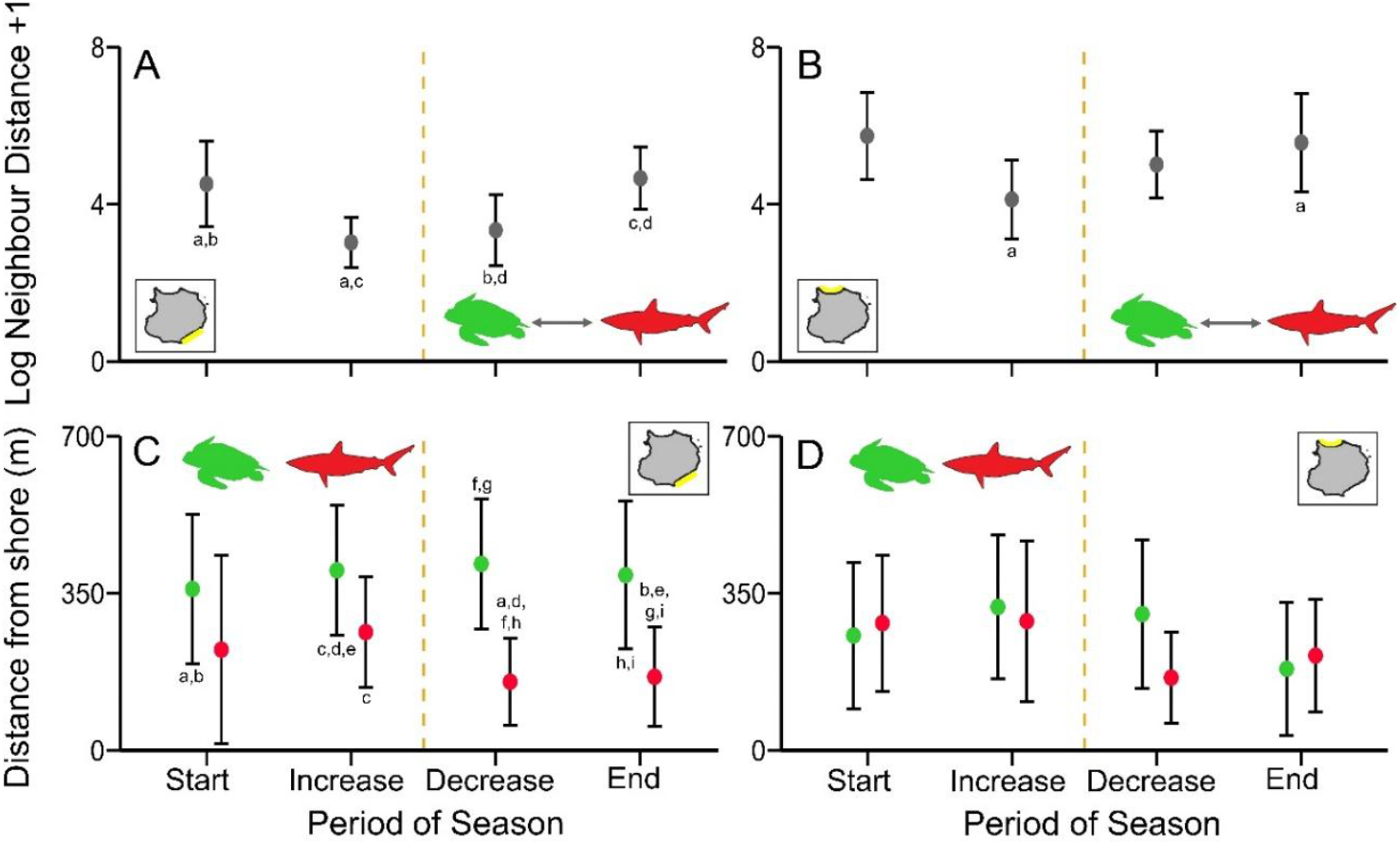
The log +1 transformed neighbour distance between sharks (red) and turtles (green) in the (**A**) high nesting density (southeast) and (**B**) low nesting density (north) areas for turtles. Distance from the coastline for sea turtles and sharks in the (**C**) high nesting density and (**D**) low nesting density turtle areas throughout the sea turtle nesting season. The yellow dashed line shows when sea turtle hatchlings likely begin to emerge from nests. Error bars give mean ± standard deviation. Matching letters show significant differences.

To test whether the community dynamics were the result of density-dependent processes, the distances between the coastline and individual sharks and turtles were calculated. The distance from the shoreline correlated with an interaction between the marine megafauna groups, in both the low- and high nesting density areas at each period of the nesting season (LMEM: F_*_= 2.40, *p* = 0.025; Figure 4c, d, Figure S4). We found in the high nesting density area, as sea turtles arrived in large numbers, they remained further from the shoreline than sharks, with sharks positioned between the turtles and the coastline (Increase period: 304.50 ± 82.51 m, *p* = 0.042; Decrease: 228.32 ± 17.03 m, *p* < 0.001; End: 224.58 ± 38.08 m, *p* <0.001; Figure 4c). No such spatial structuring was detected at the low nesting density area, neither for the sharks nor any of the other marine megafauna groups (all *p* > 0.05; Figure 4d; Figure S4b). Noteworthy, turtles in the high nesting density area were further from shore than those in the low-density area (End season: 182.88 ± 32.12 m, *p* < 0.001; Figure 4c, d). Together those results suggest a cryptic spatial organization of the different taxonomic groups in the coastal areas in front of nesting beaches.

## Discussion

Migration is a complex life history trait that impacts both individuals and communities (Dingle 2014; Shaw 2016). Here, we explored how marine community structures change throughout the mass migration of loggerhead sea turtled during their nesting season, in Cabo Verde, a marine biodiversity hotspot. As Cabo Verde holds the world’s largest nesting population of loggerhead sea turtles (Taxonera *et al.* 2022), this region is of global ecological and conservation interest. We found that in the high nesting density area, the influx of adult turtles temporarily displaces other marine megafauna species like sharks, reshaping community composition and spatial community structure. As the season progressed, spatial overlap increased between sharks and turtles in the high nesting density area. Yet, sharks predominantly remained between the shoreline and turtles, potentially indicating the predatory targeting of emerging hatchlings, an ephemeral resource pulse, rather than adult turtles. Contrastingly, in the low nesting density area, the community composition and spatial structure remained relatively unchanged, suggesting an ecological density threshold must be met for turtles to drive structural changes. As the nesting season ends, and turtles depart, the community return towards its original composition and spatial structure indicating a cyclical reshaping.

While the Cabo Verde archipelago hosts the world’s largest nesting aggregation of loggerhead turtles, their in-water abundance and interactions with other species remain largely elusive. We found turtles generally aggregated in water close to their nesting beaches in proportional densities to nesting, similar to observations in Greece, the USA and Brazil (Dickson *et al.* 2022; Hart *et al.* 2013; Marcovaldi *et al.* 2010). Remaining close to nesting areas allows turtles to minimise their movements between in-water and nesting beaches, possibly allocating energy resource into egg development (Fossette *et al.* 2012; Shaver *et al.* 2017). As expected, the strongest signals of community composition change and spatial structure were detected in the high nesting density area, with the communities in the low nesting density area showing no composition or spatial changes. This result suggests the mass influx of adult turtles triggers niche partitioning, where turtles displace other species, such as sharks within a constrained area. These density-dependent dynamics parallel invasive species ecology, whereby invaders outcompete native species (Bewick *et al.* 2017; Mooney & Cleland 2001). However, our observations show only temporary displacement. Community composition and spatial stability in the low nesting density area further reinforces the density-dependent nature of the community effects of migration. What remains unknown, however, is whether there exists a tipping point or a sea turtle density threshold inducing community spatial restructuring. Ecological thresholds however have revealed how community compositions change in response to environmental (e.g., temperature) or ecological (e.g., the influx of migratory species) changes in the ecosystem (Baker & King 2010; Groffman *et al.* 2006; Osman *et al.* 2010). For instance, thresholds being met due to aridification results in some plant species losing coverage and richness (Berdugo *et al.* 2020). The mass influx of migratory species can prompt the relocation of a resident species, so residents can limit competition for resources (Powell *et al.* 2021). As our study was not focused on when the exact threshold, if it exists, was met, and future studies should focus on the whole of Cabo Verde as many environmental and nesting density gradients exist across the archipelago to resolve this question.

By calculating shark-turtle distances throughout the nesting period, we found that sharks and turtles became physically closer as the season progressed in the high nesting density areas. However, we also showed that sharks were distributed between the shoreline and the turtles,revealing a cryptic structure within a restricted shallow area. In the high nesting density area, increased shark presence aligning with hatchling emergence (after ∼50-60 days of incubation post-nesting) points towards active foraging on emerging turtle hatchlings from the nesting beaches. Small shark species (e.g., blacktip reef (*Carcharhinus melanopterus*), smoothhound (*Mustelus mustelus*)) and juvenile sharks do consume hatchlings in Boa Vista (Bashir *et al.* 2020; Louro *et al.* 2022), whereas larger shark species, such as tiger sharks, can feed on adult turtles as seen in Australia and the USA (Hammerschlag *et al.* 2015; Heithaus *et al.* 2005). Predators migrating to maximise foraging activities in response to resource pulses has been detected across terrestrial, marine, and freshwater ecosystems (e.g., North Atlantic right whales (*Eubalaena glacialis*) to feed on copepods, Firestone *et al.* 2008; Furey *et al.* 2018).

As adult turtles were positioned further away from the shoreline compared to sharks, this may suggest they attempt at reducing predation risk or are plastic enough to use deeper resting sites. Turtles may aggregate in large numbers to adopt grouping strategies to evade predators (Hintz & Lonzarich 2018; Ioannou *et al.* 2009), enhance collective vigilance (Dröge *et al.* 2017), act as multiple targets to confuse predators (Ioannou *et al.* 2009) or enable members of the group to escape when another member is captured (Foster & Treherne 1981). There is evidence to suggest turtles use grouping strategies, as hatchlings emerge simultaneously (Santos *et al.* 2016), while adult olive ridley turtles (*Lepidochelys olivacea*) in Costa Rica perform mass nesting events (*arribada*) to possibly dilute predation risk (Eckrich & Owens 1995). Alternatively, sea turtles can exploit deeper areas for resting between nesting events, as shown by tracking turtles from Boa Vista and characterizing resting phases (Gunner *et al.* 2021) These deeper zones may offer thermal stability and fewer predators, providing a relatively safe refuge during the inter-nesting intervals.

Given the flagship nature of marine megafauna, from a conservation point of view, protecting key habitats and managing human activities through regulations is imperative for preserving ecological communities during critical periods of the year (Hooker & Gerber 2004; Lascelles *et al.* 2014). Prioritizing nearshore protection year-round and offshore protection during peak nesting would enable community dynamics to be preserved. Overall, tracking this cyclic community dynamics has highlighted critical conservation opportunities and shed light on fundamental ecological processes of niche partitioning at high density of migratory individuals.

## Supporting information

Supplementary tables and figures that support the manuscript

## Acknowledgements

This work was funded by Queen Mary University of London (UK), the National Environment Research Council (UK) and the London NERC Doctoral Training Partnership (UK). Permits were delivered by the Cabo Verde Ministry of Environment (Authorization: 037/DNA/2021 to CE). We also thank Fundação Tartaruga, Cabo Verde Natura 2000 and Bios CV for their support during the collection of fieldwork. We would like to thank Dr. Tom Fayle for their insightful comments on a previous version of the manuscript.

## Notes

### Competing Interest Statement

The authors have declared no competing interest.

## References

Anderwald, P., Herfindal, I., Haller, R.M., Risch, A.C., Schütz, M., Schweiger, A.K., et al. (2015). Influence of migratory ungulate management on competitive interactions with resident species in a protected area. Ecosphere, 6, art228.

Baker, M.E. & King, R.S. (2010). A new method for detecting and interpreting biodiversity and ecological community thresholds. Methods Ecol. Evol., 1, 25–37.

Baltazar-Soares, M., Klein, J.D., Correia, S.M., Reischig, T., Taxonera, A., Roque, S.M., et al. (2020). Distribution of genetic diversity reveals colonization patterns and philopatry of the loggerhead sea turtles across geographic scales. Sci. Rep., 10, 18001.

Bashir, Z., Abdullah, M.M., Ghaffar, M.Abd. & Rusli, M.U. (2020). Exclusive predation of sea turtle hatchlings by juvenile blacktip reef sharks Carcharhinus melanopterus at a turtle nesting site in Malaysia. J. Fish Biol., 97, 1876–1879.

Bauer, S. & Hoye, B.J. (2014). Migratory Animals Couple Biodiversity and Ecosystem Functioning Worldwide. Science, 344, 1242552.

Bellard, C., Bertelsmeier, C., Leadley, P., Thuiller, W. & Courchamp, F. (2012). Impacts of climate change on the future of biodiversity. Ecol. Lett., 15, 365–377.

Berdugo, M., Delgado-Baquerizo, M., Soliveres, S., Hernández-Clemente, R., Zhao, Y., Gaitán, J.J., et al. (2020). Global ecosystem thresholds driven by aridity. Science, 367, 787–790.

Bewick, S., Wang, G., Younes, H., Li, B. & Fagan, W.F. (2017). Invasion dynamics of competing species with stage-structure. J. Theor. Biol., 435, 12–21.

Bjorndal, K.A. & Jackson, J.B.C. (2003). Roles of sea turtles in marine ecosystems: reconstructing the past. In: The Biology of Sea Turtles. CRC Press, pp. 259–273.

Bro-Jørgensen, J., Franks, D.W. & Meise, K. (2019). Linking behaviour to dynamics of populations and communities: application of novel approaches in behavioural ecology to conservation. Philos. Trans. R. Soc. B Biol. Sci., 374, 20190008.

Brönmark, C., Hulthén, K., Nilsson, P.A., Skov, C., Hansson, L.-A., Brodersen, J., et al. (2014). There and back again: migration in freshwater fishes. Can. J. Zool., 92, 467– 479.

Calenge, C. (2006). The package “adehabitat” for the R software: A tool for the analysis of space and habitat use by animals. Ecol. Model., 197, 516–519.

Chapman, J.W., Reynolds, D.R. & Wilson, K. (2015). Long-range seasonal migration in insects: mechanisms, evolutionary drivers and ecological consequences. Ecol. Lett., 18, 287–302.

Costa-Pereira, R., Moll, R.J., Jesmer, B.R. & Jetz, W. (2022). Animal tracking moves community ecology: Opportunities and challenges. J. Anim. Ecol., 91, 1334–1344.

Dickson, L.C.D., Tugwell, H., Katselidis, K.A. & Schofield, G. (2022). Aerial Drones Reveal the Dynamic Structuring of Sea Turtle Breeding Aggregations and Minimum Survey Effort Required to Capture Climatic and Sex-Specific Effects. Front. Mar. Sci., 9.

Dingle, H. (2014). Migration: The Biology of Life on the Move. 2nd edn. Oxford University Press, Oxford.

Dröge, E., Creel, S., Becker, M.S. & M’soka, J. (2017). Risky times and risky places interact to affect prey behaviour. Nat. Ecol. Evol., 1, 1123–1128.

Duarte, M.C. & Romeiras, M. (2009). Cape Verde Islands. Encycl. Isl.

Dujon, A.M., Ierodiaconou, D., Geeson, J.J., Arnould, J.P.Y., Allan, B.M., Katselidis, K.A., et al. (2021). Machine learning to detect marine animals in UAV imagery: effect of morphology, spacing, behaviour and habitat. Remote Sens. Ecol. Conserv., 7, 341– 354.

Eckrich, C.E. & Owens, D.Wm. (1995). Solitary versus Arribada Nesting in the Olive Ridley Sea Turtles (Lepidochelys Olivacea): A Test of the Predator-Satiation Hypothesis. Herpetologica, 51, 349–354.

Firestone, J., Lyons, S.B., Wang, C. & Corbett, J.J. (2008). Statistical modeling of North Atlantic right whale migration along the mid-Atlantic region of the eastern seaboard of the United States. Biol. Conserv., 141, 221–232.

Fitzpatrick, R., Thums, M., Bell, I., Meekan, M.G., Stevens, J.D. & Barnett, A. (2012). A Comparison of the Seasonal Movements of Tiger Sharks and Green Turtles Provides Insight into Their Predator-Prey Relationship. PLoS ONE, 7, e51927.

Fossette, S., Schofield, G., Lilley, M.K.S., Gleiss, A.C. & Hays, G.C. (2012). Acceleration data reveal the energy management strategy of a marine ectotherm during reproduction. Funct. Ecol., 26, 324–333.

Foster, W.A. & Treherne, J.E. (1981). Evidence for the dilution effect in the selfish herd from fish predation on a marine insect. Nature, 293, 466–467.

Furey, N.B., Armstrong, J.B., Beauchamp, D.A. & Hinch, S.G. (2018). Migratory coupling between predators and prey. Nat. Ecol. Evol., 2, 1846–1853.

Giroux, M.-A., Berteaux, D., Lecomte, N., Gauthier, G., Szor, G. & Bêty, J. (2012). Benefiting from a migratory prey: spatio-temporal patterns in allochthonous subsidization of an arctic predator. J. Anim. Ecol., 81, 533–542.

Godley, B.J., Barbosa, C., Bruford, M., Broderick, A.C., Catry, P., Coyne, M.S., et al. (2010). Unravelling migratory connectivity in marine turtles using multiple methods: Migratory connectivity in marine turtles. J. Appl. Ecol., 47, 769–778.

Grémillet, D., Chevallier, D. & Guinet, C. (2022). Big data approaches to the spatial ecology and conservation of marine megafauna. ICES J. Mar. Sci., 79, 975–986.

Groffman, P.M., Baron, J.S., Blett, T., Gold, A.J., Goodman, I., Gunderson, L.H., et al. (2006). Ecological Thresholds: The Key to Successful Environmental Management or an Important Concept with No Practical Application? Ecosystems, 9, 1–13.

Gunner, R., Wilson, R., Holton, M., Scott, R., Arkwright, A., Fahlman, A., et al. (2021). Activity of loggerhead turtles during the U-shaped dive: insights using angular velocity metrics. Endanger. Species Res., 45, 1–12.

Hammerschlag, N., Broderick, A.C., Coker, J.W., Coyne, M.S., Dodd, M., Frick, M.G., et al. (2015). Evaluating the landscape of fear between apex predatory sharks and mobile sea turtles across a large dynamic seascape. Ecology, 96, 2117–2126.

Hart, C.E., Ley-Quiñonez, C., Maldonado-Gasca, A., Zavala, A. & Abreu-Grobois, F.A. (2014). Nesting characteristics of olive ridley turtles (Lepidochelys olivacea) on El Naranjo beach, Nayarit, Mexico. Herpetol. Conserv. Biol., 9, 524–534.

Hart, K.M., Lamont, M.M., Sartain, A.R., Fujisaki, I. & Stephens, B.S. (2013). Movements and Habitat-Use of Loggerhead Sea Turtles in the Northern Gulf of Mexico during the Reproductive Period. PLOS ONE, 8, e66921.

Hays, G.C. & Hawkes, L.A. (2018). Satellite Tracking Sea Turtles: Opportunities and Challenges to Address Key Questions. Front. Mar. Sci., 5.

Heithaus, M., Frid, A., Wirsing, A., Bejder, L. & Dill, L. (2005). Biology of sea turtles under risk from tiger sharks at a foraging ground. Mar. Ecol. Prog. Ser., 288, 285–294.

Heithaus, M.R., Wirsing, A.J., Thomson, J.A. & Burkholder, D.A. (2008). A review of lethal and non-lethal effects of predators on adult marine turtles. J. Exp. Mar. Biol. Ecol., 356, 43–51.

Hintz, W.D. & Lonzarich, D.G. (2018). Maximizing foraging success: the roles of group size, predation risk, competition, and ontogeny. Ecosphere, 9, e02456.

Hooker, S.K. & Gerber, L.R. (2004). Marine Reserves as a Tool for Ecosystem-Based Management: The Potential Importance of Megafauna. BioScience, 54, 27.

Hulbert, L.B., Aires‐da‐Silva, A.M., Gallucci, V.F. & Rice, J.S. (2005). Seasonal foraging movements and migratory patterns of female Lamna ditropis tagged in Prince William Sound, Alaska. J. Fish Biol., 67, 490–509.

Ioannou, C.C., Morrell, L.J., Ruxton, G.D. & Krause, J. (2009). The Effect of Prey Density on Predators: Conspicuousness and Attack Success Are Sensitive to Spatial Scale. Am. Nat., 173, 499–506.

Kelaher, B.P., Colefax, A.P., Tagliafico, A., Bishop, M.J., Giles, A. & Butcher, P.A. (2020). Assessing variation in assemblages of large marine fauna off ocean beaches using drones. Mar. Freshw. Res., 71, 68.

Lascelles, B., Notarbartolo Di Sciara, G., Agardy, T., Cuttelod, A., Eckert, S., Glowka, L., et al. (2014). Migratory marine species: their status, threats and conservation management needs. Aquat. Conserv. Mar. Freshw. Ecosyst., 24, 111–127.

Levi, T., Wheat, R.E., Allen, J.M. & Wilmers, C.C. (2015). Differential use of salmon by vertebrate consumers: implications for conservation. PeerJ, 3, e1157.

Louro, A., Almeida, L., Marco, A., Martins, S. & Lopes, E.P. (2022). High marine predation of loggerhead turtle hatchlings at Boavista, Cabo Verde. Zool. Caboverdiana, 10, 26– 34.

Marcovaldi, M.Â., Lopez, G.G., Soares, L.S., Lima, E.H.S.M., Thomé, J.C.A. & Almeida, A.P. (2010). Satellite-tracking of female loggerhead turtles highlights fidelity behavior in northeastern Brazil. Endanger. Species Res., 12, 263–272.

Mooney, H.A. & Cleland, E.E. (2001). The evolutionary impact of invasive species. Proc.Natl. Acad. Sci., 98, 5446–5451.

Oksanen, J., Simpson, G.L., Blanchet, F.G., Kindt, R., Legendre, P., Minchin, P.R., et al.(2022). Vegan: Community Ecology Package.

Osman, R.W., Munguia, P. & Zajac, R.N. (2010). Ecological thresholds in marine communities: theory, experiments and management. Mar. Ecol. Prog. Ser., 413, 185–187.

Ostfeld, R.S. & Keesing, F. (2000). Pulsed resources and community dynamics of consumers in terrestrial ecosystems. Trends Ecol. Evol., 15, 232–237.

Powell, L.L., Ames, E.M., Wright, J.R., Matthiopoulos, J. & Marra, P.P. (2021). Interspecific competition between resident and wintering birds: experimental evidence and consequences of coexistence. Ecology, 102, e03208.

R Core Team. (2022). R: A language and environment for statistical computing.

Rosa, R., Nunes, E., Pissarra, V., Santos, C.P., Varela, J., Baptista, M., et al. (2023). Evidence for the first multi-species shark nursery area in Atlantic Africa (Boa Vista Island, Cabo Verde). Front. Mar. Sci., 10.

Santos, R.G., Pinheiro, H.T., Martins, A.S., Riul, P., Bruno, S.C., Janzen, F.J., et al. (2016). The anti-predator role of within-nest emergence synchrony in sea turtle hatchlings. Proc. R. Soc. B Biol. Sci., 283, 20160697.

Schlägel, U.E., Grimm, V., Blaum, N., Colangeli, P., Dammhahn, M., Eccard, J.A., et al. (2020). Movement-mediated community assembly and coexistence. Biol. Rev., 95, 1073–1096.

Schofield, G., Papafitsoros, K., Chapman, C., Shah, A., Westover, L., Dickson, L.C.D., et al. (2022). More aggressive sea turtles win fights over foraging resources independent of body size and years of presence. Anim. Behav., 190, 209–219.

Shaver, D.J., Hart, K.M., Fujisaki, I., Bucklin, D., Iverson, A.R., Rubio, C., et al. (2017). Internesting movements and habitat-use of adult female Kemp’s ridley turtles in the Gulf of Mexico. PLOS ONE, 12, e0174248.

Shaw, A.K. (2016). Drivers of animal migration and implications in changing environments. Evol. Ecol., 30, 991–1007.

Stokes, H.J., Mortimer, J.A., Laloë, J.-O., Hays, G.C. & Esteban, N. (2023). Synergistic use of UAV surveys, satellite tracking data, and mark-recapture to estimate abundance of elusive species. Ecosphere, 14, e4444.

Taxonera, A., Fairweather, K., Jesus, A., Gonzalves, A., Quelruga, A., Lima, A., et al. (2022). Cabo Verde: Sea Turtles “In Abundance.” SWOT State Worlds Sea Turt. Rep., 17.

Wenzel, F., Broms, F., Lopez-Suarez, P., Freire Lopes, K., Veiga, N., Yeoman, K., et al. (2020). Humpback Whales (Megaptera novaeangliae) in the Cape Verde Islands: Migratory Patterns, Resightings, and Abundance. Aquat. Mamm., 46, 21–31.

