## Supplementary tables and figures that support the manuscript for "The migration of an expanding sea turtle population alters the structure of marine megafauna communities"

**Table S1.** Species and their number of records detected in aerial drone surveys flown during the loggerhead sea turtle nesting season, with their current IUCN Red List Status. Asterisks (*) give the IUCN status of the regional population.

| Taxa | Species name | IUCN status | Number of records |
|---|---|---|---|
| ***Elasmobranch*** | | | |
| Lemon Shark | *Negaprion brevirostris* | *Vulnerable* | 52 |
| Tiger Shark | *Galeocerdo cuvier* | *Near threatened* | 28 |
| Atlantic Nurse Shark | *Ginglymostoma cirratum* | *Vulnerable* | 18 |
| Bull Shark | *Carcharhinus leucas* | *Vulnerable* | 4 |
| Devil Ray | *Mobula tarapacana* | *Endangered* | 3 |
| Blackchin Guitarfish | *Rhinobatos cemiculus* | *Critically Endangered* | 2 |
| Giant Oceanic Manta Ray | *Mobula birostris* | *Endangered* | 2 |
| Unknown Hammerhead Shark Species | *Sphyrnidae spp.* | | 4 |
| Unknown Shark Species | | | 257 |
| Unknown Mobula Ray Species | *Mobulidae spp.* | | 39 |
| Unknown Stingray Species | *Dasaytidae spp.* | | 65 |
| **Total** | | | **376** |
| ***Sea turtle*** | | | |
| Loggerhead Turtle | *Caretta caretta* | *Endangered** | 10125 |
| Green Turtle | *Chelonia mydas* | *Endangered* | 114 |
| Hawksbill Turtle | *Eretmochelys imbricata* | *Critically Endangered* | 13 |
| Olive Ridley Turtle | *Lepidochelys olivacea* | *Vulnerable* | 2 |
| Unknown Turtle Species | | | 4414 |
| **Total** | | | **14668** |
| ***Fish*** | | | |
| Ocean Sunfish | *Mola mola* | *Vulnerable* | 3 |
| Wahoo | *Acanthocybium solandri* | *Least Concern* | 1 |
| Unknown Large Fish Species | | | 167 |
| Unknown Shoaling Fish | | | 62 |
| Unknown Barracuda Species | *Sphyraenidae spp.* | | 2 |
| Unknown Small Fish Species | | | 2 |
| **Total** | | | **234** |
| **Grand Total** | | | **15278** |

**Table S2.** SIMPER analysis of the marine megafauna groups detected in the southeast of Boa Vista, at each stage of the loggerhead sea turtle nesting season. Dissimilarity given as Bray-Curtis Dissimilarity matrices, log+1 transformed. Significant contributors ($p < 0.05$) are indicated with an asterisk (*).

| Southeast Boa Vista | | | | | |
|---|---|---|---|---|---|
| | Log Mean Abundance | | | | |
| | **Start** | **Increase** | **Dissimilarity (%)** | **Significance** | |
| Sea Turtle | 2.414 | 6.061 | 64.1 | <0.001 | * |
| Coastal Ray | 0 | 0.664 | 11 | 0.002 | * |
| Individual Fish | 0.381 | 0.649 | 10.9 | 0.813 | |
| Shark | 0.315 | 0.414 | 8.3 | 1 | |
| Fish Shoal | 0.108 | 0.207 | 4.8 | 0.538 | |
| Oceanic Ray | 0 | 0.058 | 0.9 | 0.753 | |
| | **Start** | **Decrease** | | | |
| Sea Turtle | 2.414 | 5.794 | 57 | 0.001 | * |
| Shark | 0.315 | 1.079 | 14.4 | 0.952 | |
| Coastal Ray | 0 | 0.918 | 14 | <0.001 | * |
| Individual Fish | 0.381 | 0.289 | 7.2 | 0.997 | |
| Oceanic Ray | 0 | 0.373 | 3.8 | 0.053 | |
| Fish Shoal | 0.108 | 0.207 | 3.6 | 0.745 | |
| | **Start** | **End** | | | |
| Sea Turtle | 2.414 | 2.951 | 53.8 | 0.552 | |
| Shark | 0.315 | 0.35 | 16.9 | 0.998 | |
| Individual Fish | 0.381 | 0.161 | 15.8 | 0.657 | |
| Coastal Ray | 0 | 0.199 | 6 | 0.929 | |
| Fish Shoal | 0.108 | 0.077 | 5.6 | 0.661 | |
| Oceanic Ray | 0 | 0.077 | 1.9 | 0.616 | |
| | **Increase** | **Decrease** | | | |
| Shark | 0.414 | 1.079 | 24.8 | 0.998 | |
| Sea Turtle | 6.061 | 5.794 | 23.6 | 1 | |
| Coastal Ray | 0.664 | 0.918 | 18.9 | 0.167 | |
| Individual Fish | 0.649 | 0.289 | 16.6 | 0.993 | |
| Oceanic Ray | 0.058 | 0.373 | 8.1 | 0.111 | |
| Fish Shoal | 0.207 | 0.207 | 8 | 0.741 | |
| | **Increase** | **End** | | | |
| Sea Turtle | 6.061 | 2.951 | 61.2 | 0.006 | * |
| Individual Fish | 0.649 | 0.161 | 11.7 | 0.878 | |
| Coastal Ray | 0.664 | 0.199 | 11.4 | 0.011 | * |
| Shark | 0.414 | 0.35 | 9.6 | 1 | |
| Fish Shoal | 0.207 | 0.077 | 4.2 | 0.735 | |
| Oceanic Ray | 0.058 | 0.077 | 1.9 | 0.491 | |
| | **Decrease** | **End** | | | |
| Sea Turtle | 5.794 | 2.951 | 54.9 | 0.064 | |
| Shark | 1.079 | 0.35 | 16.9 | 0.929 | |
| Coastal Ray | 0.918 | 0.199 | 14.4 | <0.001 | * |
| Individual Fish | 0.289 | 0.161 | 5.7 | 1 | |
| Oceanic Ray | 0.373 | 0.077 | 5 | 0.018 | * |
| Fish Shoal | 0.207 | 0.077 | 3.1 | 0.88 | |

**Table S3.** SIMPER analysis of the marine megafauna groups detected in the north of Boa Vista, at each stage in the loggerhead sea turtle nesting season. Dissimilarity given as Bray-Curtis Dissimilarity matrices, log+1 transformed. Significant contributors ($p$ < 0.05) are indicated with an asterisk (*).

| North Boa Vista | | | | | |
|---|---|---|---|---|---|
| | Log Mean Abundance | | | | |
| | **Start** | **Increase** | **Dissimilarity (%)** | **Significance** | |
| Sea Turtle | 1.873 | 4.278 | 51.7 | 0.073 | |
| Shark | 0.862 | 0.921 | 17.4 | 0.783 | |
| Individual Fish | 0.751 | 0.403 | 15 | 0.209 | |
| Fish Shoal | 0.147 | 0.39 | 8.4 | 0.082 | |
| Coastal Ray | 0.061 | 0.323 | 5.6 | 0.782 | |
| Oceanic Ray | 0.116 | 0 | 1.9 | 0.479 | |
| | **Start** | **Decrease** | | | |
| Sea Turtle | 1.873 | 2.784 | 35.5 | 1 | |
| Shark | 0.862 | 1.509 | 33.1 | 0.003 | * |
| Individual Fish | 0.751 | 0.655 | 21 | 0.086 | |
| Fish Shoal | 0.147 | 0 | 4 | 0.858 | |
| Coastal Ray | 0.061 | 0.116 | 3.7 | 0.992 | |
| Oceanic Ray | 0.116 | 0 | 2.7 | 0.445 | |
| | **Start** | **End** | | | |
| Sea Turtle | 1.873 | 1.62 | 32 | 0.999 | |
| Shark | 0.862 | 1.127 | 28.6 | 0.002 | * |
| Individual Fish | 0.751 | 0.6 | 24.8 | <0.001 | * |
| Fish Shoal | 0.147 | 0.258 | 9.2 | 0.066 | |
| Coastal Ray | 0.061 | 0.092 | 2.8 | 0.997 | |
| Oceanic Ray | 0.116 | 0 | 2.6 | 0.327 | |
| | **Increase** | **Decrease** | | | |
| Sea Turtle | 4.278 | 2.784 | 40.2 | 0.996 | |
| Shark | 0.921 | 1.509 | 28.3 | 0.251 | |
| Individual Fish | 0.403 | 0.655 | 15.7 | 0.771 | |
| Fish Shoal | 0.39 | 0 | 8.5 | 0.344 | |
| Coastal Ray | 0.323 | 0.116 | 7.3 | 0.795 | |
| Oceanic Ray | 0 | 0 | 0 | 0.843 | |
| | **Increase** | **End** | | | |
| Sea Turtle | 4.278 | 1.62 | 57 | 0.001 | * |
| Shark | 0.921 | 1.127 | 18.5 | 0.339 | |
| Individual Fish | 0.403 | 0.6 | 11.5 | 0.657 | |
| Fish Shoal | 0.39 | 0.258 | 7.8 | 0.089 | |
| Coastal Ray | 0.323 | 0.092 | 5.3 | 0.739 | |
| Oceanic Ray | 0 | 0 | 0 | 0.886 | |
| | **Decrease** | **End** | | | |
| Sea Turtle | 2.784 | 1.62 | 41.1 | 0.916 | |
| Shark | 1.509 | 1.127 | 30.8 | 0.001 | * |
| Individual Fish | 0.655 | 0.6 | 19 | 0.053 | |
| Fish Shoal | 0 | 0.258 | 5.7 | 0.519 | |
| Coastal Ray | 0.116 | 0.092 | 3.5 | 0.982 | |
| Oceanic Ray | 0 | 0 | 0 | 0.884 | |

**Table S4.** SIMPER analysis of the marine megafauna groups detected between the north and southeast of Boa Vista, at each stage in the loggerhead sea turtle nesting season. Dissimilarity given as Bray-Curtis Dissimilarity matrices, log+1 transformed. Significant contributors ($p$ < 0.05) are indicated with an asterisk (*).

| | Log Mean Abundance | | | | |
|---|---|---|---|---|---|
| **Start of the Season** | **Southeast** | **North** | **Dissimilarity (%)** | **Significance** | |
| Sea Turtle | 2.414 | 1.873 | 41.1 | 0.959 | |
| Shark | 0.315 | 0.862 | 24.2 | 0.129 | |
| Individual Fish | 0.381 | 0.751 | 23.3 | <0.001 | * |
| Fish Shoal | 0.108 | 0.147 | 7.3 | 0.286 | |
| Oceanic Ray | 0 | 0.116 | 2.8 | 0.305 | |
| Coastal Ray | 0 | 0.061 | 1.3 | 1 | |
| **Increase Period** | **Southeast** | **North** | **Dissimilarity (%)** | **Significance** | |
| Sea Turtle | 6.061 | 4.278 | 41.7 | 1 | |
| Shark | 0.414 | 0.921 | 18.3 | 0.997 | |
| Individual Fish | 0.649 | 0.403 | 15.4 | 0.947 | |
| Coastal Ray | 0.664 | 0.323 | 14 | 0.165 | |
| Fish Shoal | 0.207 | 0.39 | 9.3 | 0.402 | |
| Oceanic Ray | 0.058 | 0 | 1.3 | 0.855 | |
| **Decrease Period** | **Southeast** | **North** | **Dissimilarity (%)** | **Significance** | |
| Sea Turtle | 5.794 | 2.784 | 51 | 0.271 | |
| Shark | 1.079 | 1.509 | 19.7 | 0.649 | |
| Coastal Ray | 0.918 | 0.116 | 13.6 | <0.001 | * |
| Individual Fish | 0.289 | 0.655 | 9.4 | 0.979 | |
| Oceanic Ray | 0.373 | 0 | 4 | 0.113 | |
| Fish Shoal | 0.207 | 0 | 2.2 | 0.933 | |
| **End of the Season** | **Southeast** | **North** | **Dissimilarity (%)** | **Significance** | |
| Sea Turtle | 2.951 | 1.62 | 46.7 | 0.143 | |
| Shark | 0.35 | 1.127 | 26.5 | <0.001 | * |
| Individual Fish | 0.161 | 0.6 | 13.6 | 0.326 | |
| Fish Shoal | 0.077 | 0.258 | 6.3 | 0.269 | |
| Coastal Ray | 0.199 | 0.092 | 5.5 | 0.792 | |
| Oceanic Ray | 0.077 | 0 | 1.4 | 0.627 | |

### Supplementary Figures

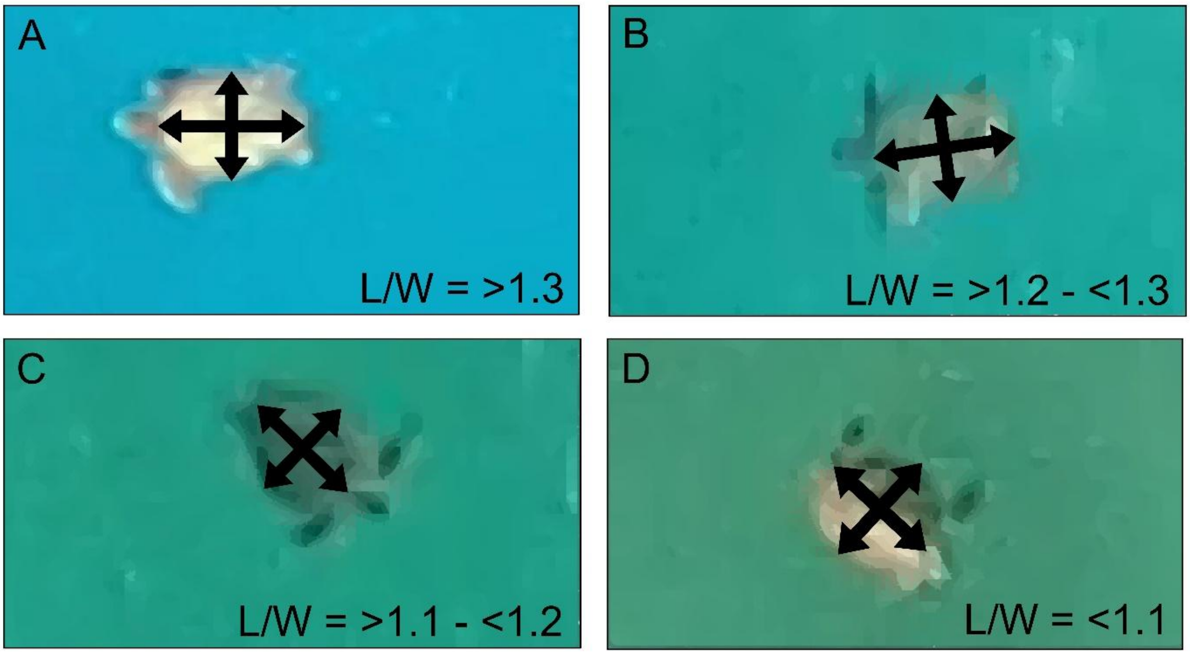


**Figure S1.** Length/width (L/W) ratios of (**A**) loggerhead sea turtles, (**B**) hawksbill turtles, (**C**) green turtles and (**D**) olive ridley turtles. Loggerhead sea turtle L/W ratios are derived from curved carapace length and width measures (Baltazar-Soares *et al.*, 2020). Green and hawksbill turtle L/W ratios are derived from straight carapace length and width (Stokes *et al.* 2023). Olive ridley turtles' L/W ratios are derived from (Hart *et al.* 2014). Image magnification at 11x to measure straight carapace length and width of turtles.

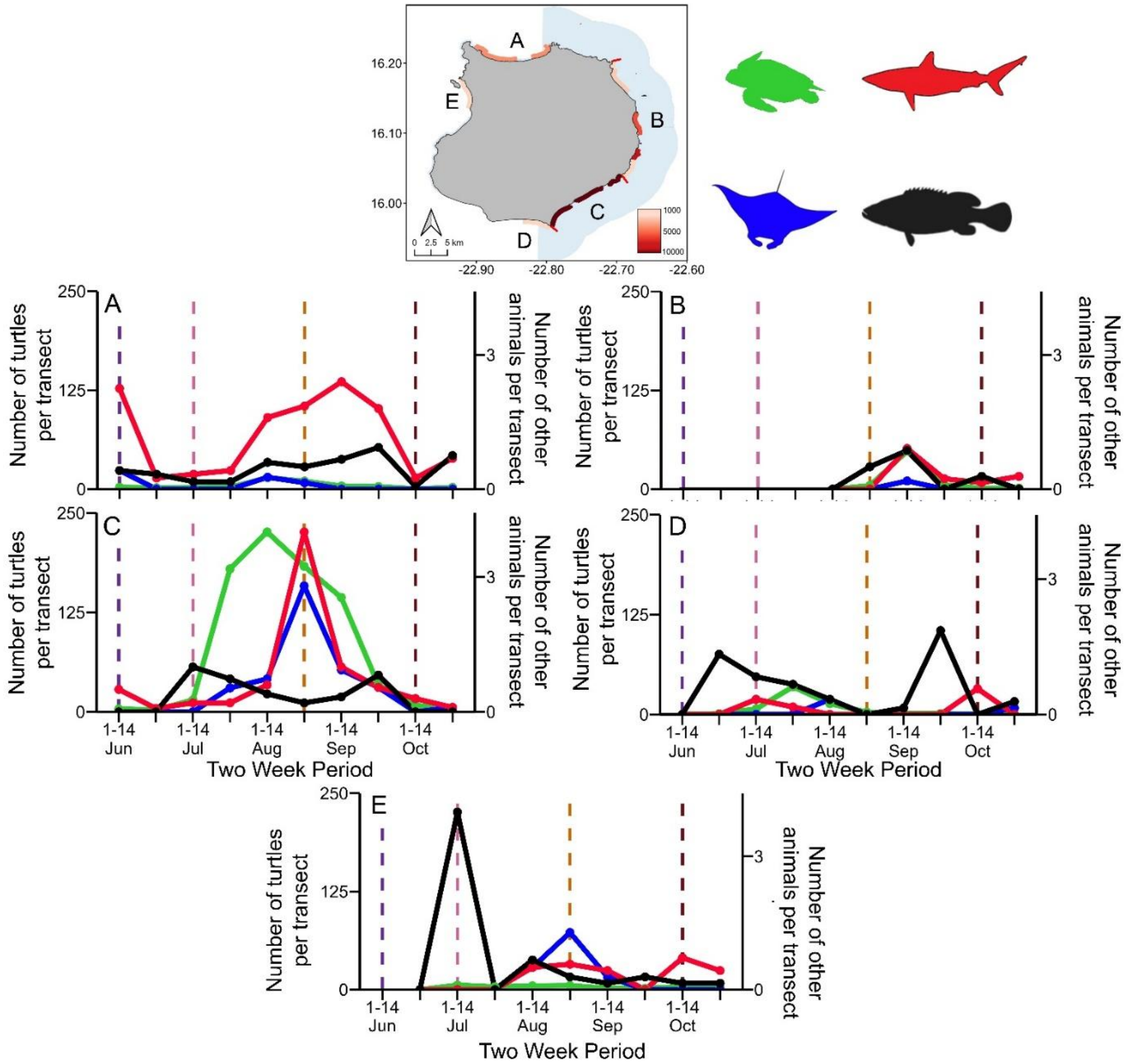


**Figure S2.** Number of sea turtles (green), sharks (red), rays (blue) and fish (black) per transect survey throughout the loggerhead sea turtle nesting season in the (A) north, (B) east, (C) southeast, (D) south and (E) west of Boa Vista, Cabo Verde. Dashed lines indicate the start of a stage in the nesting season derived from sea turtle trends; start of the season (purple), turtle population increase (pink), turtle population decrease (orange) and the end of the season (brown) periods.

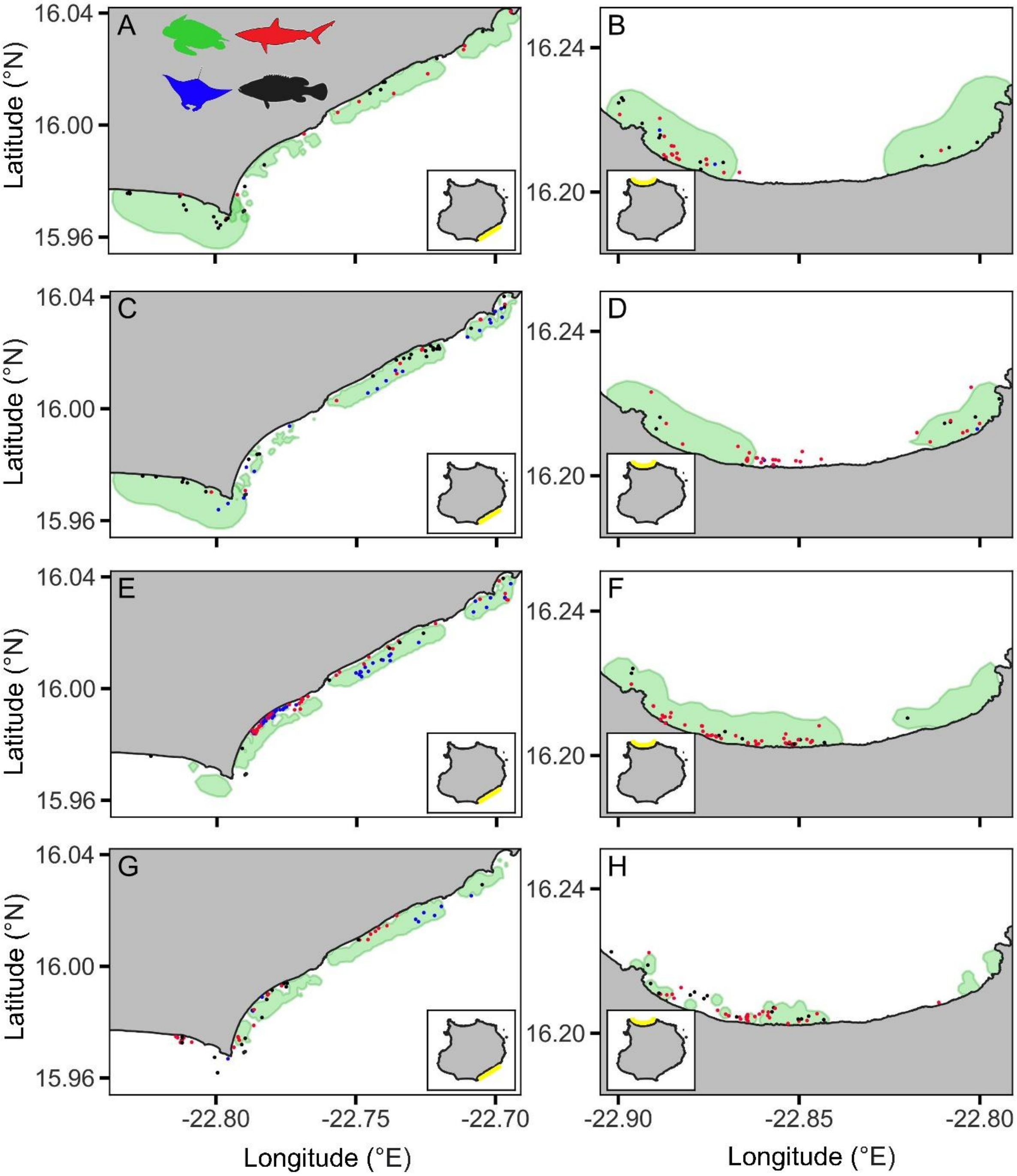


**Figure S3.** Sea turtle (green polygons) space use in the southeast (left side) and north (right side) of Boa Vista, Cabo Verde throughout the nesting season (**A-B**: start of nesting season, **C-D**: turtle number increase period, **E-F**: turtle number decrease period and **G-H**: end of nesting season periods) in relation to shark (red points), ray (blue) and fish (black) distributions.

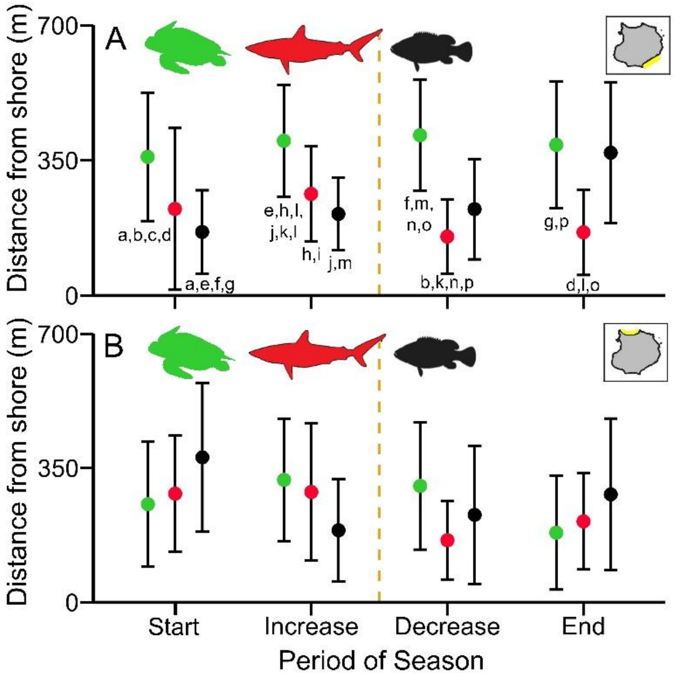


**Figure S4.** Distance from the shoreline (metres) for sea turtles (green), sharks (red) and individual fish (black) throughout the nesting season in the (**A**) southeast and (**B**) north of Boa Vista, Cabo Verde. The dashed orange line indicates the start of sea turtle hatchling emergence from nests. Points and error bars indicate mean ± standard deviation. Matching pairs of letters inside each figure give significant differences ($p < 0.05$) between means.

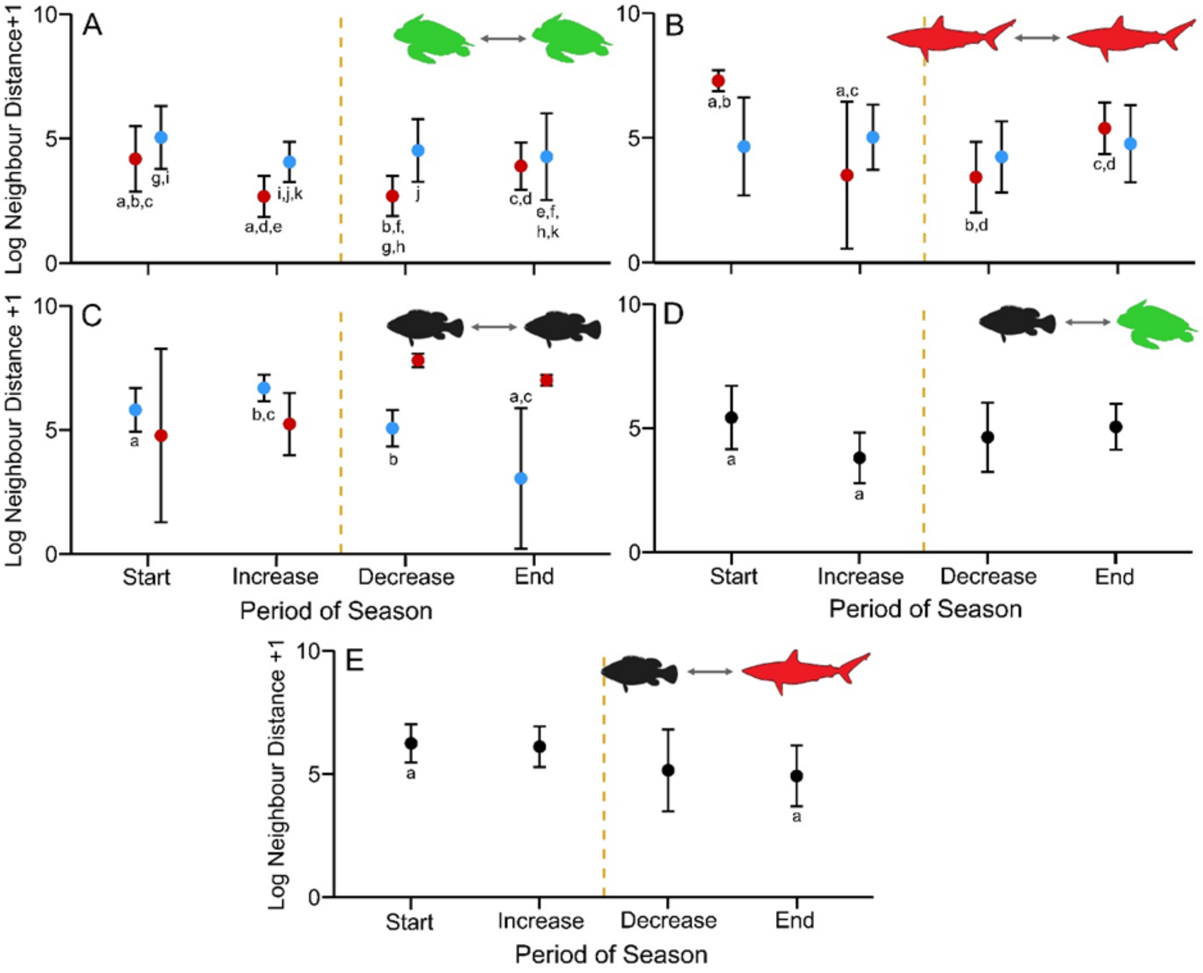


**Figure S5.** Log+1 transformed neighbour distance between (**A**) sea turtles, (**B**) between sharks and (**C**) between individual fish in the southeast (red points) and north (blue) of Boa Vista throughout the loggerhead sea turtle nesting season. (**D**) and (**E**) give log+1 transformed neighbour distance between individual fish and sea turtles, and individual fish and sharks respectively, throughout the loggerhead sea turtle nesting season. The dashed orange line indicates the start of sea turtle hatchling emergence from nests. Points and error bars indicate mean ± standard deviation. Matching pairs of letters inside each figure give significant differences ($p$ < 0.05) between means.

As expected from different nesting densities, the distance among turtles was lowest in the high-density area compared to the low-density area, which is a pattern that remained through the nesting season (LMEM: $F_{3,12766}$ = 50.77, $p$ < 0.001; Figure S5a). This result suggests that our metrics capture the dynamics and structure of the community. The distance between sharks was only different in the high-density area (LMEM: $F_{3,75}$ = 8.42; $p$ < 0.001; Figure S5b), where sharks were further away from each other at the start of the nesting season (5.01 ± 0.85, N = 6) than they were at the increase and decrease periods (increase: 3.51 ± 2.95, n = 11; decrease: 3.42 ± 1.41, n = 74; both $p < 0.001$; Figure S5b). At the end of the season (4.77 ± 1.03, n = 13) sharks were further away from each other than they were at the increase and decrease periods (increase-end: $p = 0.013$; decrease to end: $p = 0.010$; Figure C S5b).

The distance between large fish had only changed in the north throughout the nesting season (LMEM: $F_{3,92} = 10.04$, $p < 0.001$; Figure S5c), where they were further away from each other at the start of the season (5.81 ± 0.88, N = 17) than they were at the end of the season (3.04 ± 2.84, N = 26, $p = 0.001$; Figure S5c). During the increase period (6.68 ± 0.54, N = 8), fish were further away from each other, than they were at the decrease period (5.06 ± 0.73, N = 11, $p = 0.012$; Figure S5c) and the end of the season ($p < 0.001$; Figure S5c).

Between the fish and sea turtles, the distance had only changed during the nesting season regardless of the region (LMEM: $F_{3,87} = 3.83$; Figure S5d). However, fish and turtles were only further away from each other at the start of the season (5.42 ± 1.28, N = 27), than they were at the increase period (3.80 ± 1.01 N = 20) ($p = 0.010$; Figure S5d).

Like the fish-turtle distances, fish-to-shark distances only changed throughout the nesting season, regardless of the region (LMEM: $F_{3,87} = 4.68$; $p = 0.005$; Figure S5e). Here, fish and sharks were further away from each other at the start of the season (6.24 ± 0.77, N = 17) than they were at the end of the season (4.92 ± 1.24, N = 28) ($p = 0.006$; Figure S5e).
